# The Atad5 RFC-like complex is the major unloader of proliferating cell nuclear antigen in *Xenopus* egg extracts

**DOI:** 10.1101/2023.08.28.555229

**Authors:** Yoshitaka Kawasoe, Sakiko Shimokawa, Peter J. Gillespie, J. Julian Blow, Toshiki Tsurimoto, Tatsuro S. Takahashi

## Abstract

Proliferating cell nuclear antigen (PCNA) is a homo-trimeric clamp complex that serves as the molecular hub for various DNA transactions, including DNA synthesis and post-replicative mismatch repair. Its timely loading and unloading are critical for genome stability. PCNA loading is catalyzed by Replication factor C (RFC) and the Ctf18 RFC-like complex (Ctf18-RLC), and its unloading is catalyzed by Atad5/Elg1-RLC. However, RFC, Ctf18-RLC, and even some subcomplexes of their shared subunits are capable of unloading PCNA *in vitro*, leaving an ambiguity in the division of labor in eukaryotic clamp dynamics. By using a system that specifically detects PCNA unloading, we show here that Atad5-RLC, which accounts for only approximately 3% of RFC/RLCs, nevertheless provides the major PCNA unloading activity in *Xenopus* egg extracts. RFC and Ctf18-RLC each account for approximately 40% of RFC/RLCs, while immunodepletion of neither Rfc1 nor Ctf18 detectably affects the rate of PCNA unloading in our system. PCNA unloading is dependent on the ATP-binding motif of Atad5, independent of nicks on DNA and chromatin assembly, and inhibited effectively by PCNA-interacting peptides. These results support a model in which Atad5-RLC preferentially unloads DNA-bound PCNA molecules that are free from their interactors.

## Introduction

Eukaryotic chromosome replication is a complex process involving DNA synthesis, Okazaki fragment maturation, chromatin assembly, and post-replicative mismatch repair (MMR) (1, 2). These processes are tightly regulated to occur in the right order at the right place. The replication clamp, a ring-shaped protein complex structurally conserved among three domains of life, encircles DNA, translocates along DNA, and interacts with numerous factors related to replication and repair (3, 4). Its loading and unloading are critical steps for the sequential and temporal regulation of the assembly and disassembly of multiprotein complexes required for various DNA transactions.

The eukaryotic replication clamp is proliferating cell nuclear antigen (PCNA), a homo-trimeric complex whose monomer consists of two structurally similar N- and C-globular domains connected by the interdomain connecting loop. The interaction between PCNA and its binding partners is frequently mediated through a short peptide that binds to a hydrophobic pocket formed underneath the interdomain connecting loop. A major class of such short peptides is the PCNA-interacting peptide (PIP) box (5, 6), an octapeptide motif found in many PCNA interactors, including DNA polymerases, Flap structure-specific endonuclease 1 (Fen1), DNA ligase 1 (Lig1), Chromatin assembly factor 1 (CAF-1), and the MutSα (Msh2–Msh6) and MutSβ (Msh2–Msh3) mismatch sensor complexes (3, 4). Through the interaction with its binding partners, PCNA coordinates multiple reactions of chromatin replication. During lagging strand synthesis, for example, PCNA recruits DNA polymerase δ (Pol δ) onto short primers synthesized by DNA polymerase α (Pol α) to extend Okazaki fragments, Fen1 and Lig1 to process and ligate Okazaki fragments, and CAF-1 to assemble chromatin on the nascent DNA (1, 3, 4). During and after DNA synthesis, PCNA also plays multiple critical roles in MMR: it recruits MutSα and MutSβ mismatch sensors to the replication factory, activates the MutLα nicking endonuclease in a strand-specific manner, thereby functioning as a marker for the newly-synthesized DNA strand, and supports repair DNA synthesis by Pol δ (7–18). Eukaryotes also have another structurally related clamp complex, Rad9–Hus1–Rad1 (9–1–1), a hetero-trimeric complex dedicated to DNA damage checkpoint activation and repair (19).

Since free PCNA forms a closed ring structure, its DNA loading must involve the opening of the ring. This process is catalyzed by the evolutionarily conserved clamp loader complexes (19–22). The major eukaryotic clamp loader, Replication Factor C (RFC), is a hetero-pentameric complex of Rfc1–5, all of which share the AAA+ ATPase domain and are arranged in a spiral conformation (23, 24). RFC has four active ATP-binding sites, and a series of studies demonstrate that sequential ATP binding to RFC induces RFC–PCNA complex formation, the spring-washer-like opening of the PCNA ring, and the binding of the RFC–PCNA complex onto the 3′-terminus of a primer-template junction (24–27). Subsequent hydrolysis of ATP molecules facilitates the closure of the PCNA ring and the dissociation of RFC from PCNA (28).

Eukaryotes have three Rfc1 paralogs: Ctf18, Elg1 (ATAD5 in humans and Atad5 in *Xenopus*), and Rad17 (29–34). These paralogs replace the largest Rfc1 subunit of RFC to form alternative RFC-like complexes (RLCs). Two additional subunits, Ctf8 and Dcc1, are also included in Ctf18-RLC (30, 35). Unlike other RFC/RLCs, Rad17-RLC is specialized for the loading of the 9–1–1 checkpoint clamp onto the single-stranded DNA region at the 5′-terminus of a junction between single-stranded and double-stranded DNA (36–38). Ctf18-RLC has a PCNA-loading activity and forms an active PCNA-loading complex with the major leading-strand replicase DNA polymerase ε (Pol ε), and thus is proposed to be an alternative PCNA loader for the leading strand (39–43). Ctf18-RLC also promotes the establishment of sister chromatid cohesion (29, 30, 42). In contrast to RFC and Ctf18-RLC, no PCNA loading activity has been reported for Elg1/ATAD5-RLC. Rather, this complex is proposed to be a PCNA unloader, since deletion of *elg1* or depletion of its mammalian homolog ATAD5 leads to the accumulation of chromatin-bound PCNA, and purified Elg1-RLC and ATAD5-RLC exhibit the PCNA unloading activity (44–47). Deficiency of the *elg1*/*Atad5* gene in yeast and mice leads to chromosome instability, whilst also predisposing mice to cancer, suggesting that the timely unloading of PCNA is critical for genome stability (31–33, 48, 49). The regulation of PCNA unloading is also vital to MMR: MutSα interacts with PCNA and prevents its unloading to retain MMR capability (9, 13), while conversely, over-accumulation of PCNA on chromatin interferes with MMR (50).

Although evidence that Elg1/ATAD5-RLC is a primary PCNA unloader *in vivo* has been accumulated, the PCNA unloading activity is clearly not specific to this complex. RFC is a pentameric complex of Rfc1, Rfc4, Rfc3, Rfc2, and Rfc5 arranged in this order, and Rfc1 is not essential for the opening of PCNA (51). Consistently, the human RFC4–RFC3–RFC2, the yeast Rfc4–Rfc3–Rfc2–Rfc5, and even the yeast Rfc2–Rfc5 subcomplexes are capable of unloading PCNA from nicked circular DNA, suggesting that the intrinsic PCNA unloading activity resides in the smaller subunits shared by all the clamp-loader/unloader complexes (51, 52). In agreement with this notion, the PCNA unloading activity has been detected in all RFC and RLCs. In yeast, both RFC and Rad24-RLC (the yeast counterpart of Rad17-RLC) unload PCNA from singly-nicked circular DNA *in vitro* (51), and human RFC unloads PCNA from nicked circular DNA in a purified system and covalently closed DNA after replication by the SV40 system (53, 54). Likewise, yeast Ctf18-RLC unloads PCNA *in vitro* in the presence of replication protein A (RPA), a eukaryotic single-stranded DNA binding protein (55). These biochemical data raise a possibility that not only Elg1/ATAD5-RLC but also RFC, Ctf18-RLC, and Rad17-RLC contribute to PCNA unloading *in vivo*. However, the division of labor of RFC and other RLCs in PCNA unloading has not been fully understood, partly because the inactivation of clamp loaders masks their possible contribution to PCNA unloading by inhibiting the PCNA loading reaction.

To clarify the contribution of clamp loaders in PCNA unloading, we herein used an assay that specifically detects PCNA unloading in the extract of *Xenopus* eggs, a physiological *in vitro* model system that recapitulates various DNA transactions, including PCNA loading/unloading and mismatch repair. Our data pinpoint Atad5-RLC as the major PCNA unloader in *Xenopus* egg extracts and support a model in which PCNA molecules that are free from their binding partners are preferentially targeted to be unloaded by Atad5-RLC.

## Results

The nucleoplasmic extract (NPE) of *Xenopus* eggs is a nearly undiluted extract of nuclear proteins and functionally recapitulates various DNA transactions, including DNA replication, PCNA loading and unloading, and mismatch repair (13, 56, 57). To gain insight into the division of labor of clamp loaders, we first quantified the concentration of RFC and RLCs in NPE. Immunoprecipitation (IP) of a shared subunit Rfc3 removed most of the Rfc1, Ctf18, Atad5, and Rad17 proteins from NPE, suggesting that the majority of the large subunits are included in Rfc3-containing complexes (Fig. 1*A*, lane 2). Consistently, simultaneous IP of Rfc1, Ctf18, Atad5, and Rad17 co-depleted more than 90% of Rfc3 and Rfc5 from NPE, suggesting that most Rfc3 and Rfc5 are included in the RFC/RLC complexes (Fig. S1*A*). Therefore, we reasoned that the large and small RFC/RLC subunits form stoichiometric complexes in NPE. Using recombinant His-tagged Rfc3 purified from bacteria, we quantified the amount of Rfc3 in the Rfc1-IP fraction, and based on this number, we then estimated the concentration of RFC in NPE at approximately 1.1 μM. Similarly, we estimated the concentrations of Ctf18-RLC, Atad5-RLC, and Rad17-RLC at 1.3, 0.10, and 0.46 μM, respectively (Figs. 1 and S1*B*).

**Figure 1:**
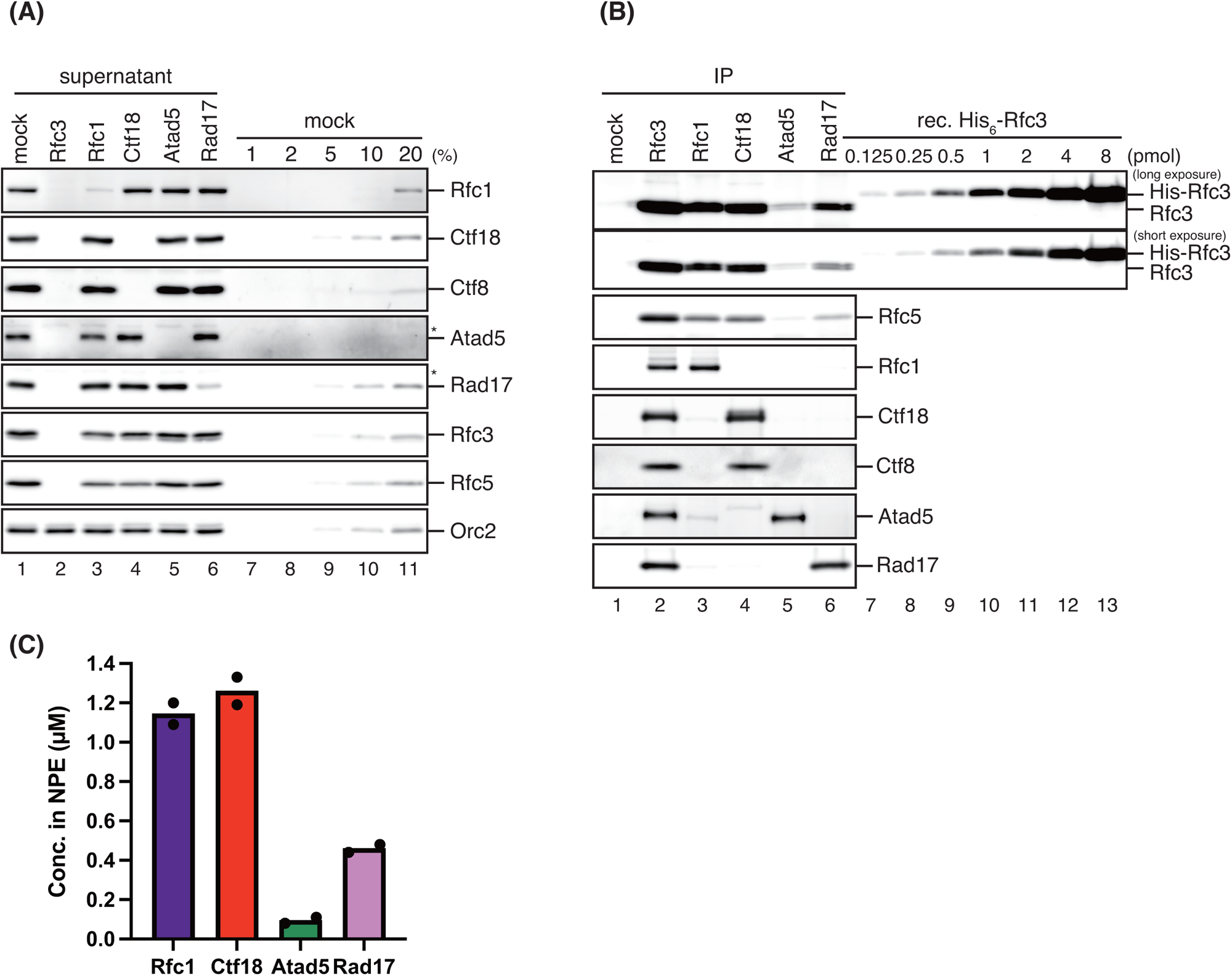
Quantification of the concentration of RFC and RLCs in NPE. (*A*, *B*) Immunoprecipitation was performed from NPE using preimmune-(“mock”, lane 1), Rfc3-(lane 2), Rfc1-(lane 3), Ctf18-(lane 4), Atad5-(lane 5), or Rad17-antibodies (lane 6), and 0.25 μL each of the supernatant (*A*) and the IP fractions corresponding to 2.5 μL each of NPE (*B*) were separated by SDS-PAGE alongside a serial dilution series of the supernatant of mock-IP NPE (*A*) or the indicated amounts of recombinant His_6_-Rfc3 (*B*), followed by immunoblotting with indicated antibodies. Orc2 serves as a loading control. (*) Cross-reacting band. (*C*) The estimated concentrations of RFC and RLCs in NPE were plotted on a graph. Filled circles represent individual data from two experiments, including the one shown in (*B*).

Biochemical studies with purified proteins have shown that not only Elg1/ATAD5-RLC but also RFC, Rad17-RLC, Ctf18-RLC, and several Rfc2–5 subcomplexes can unload PCNA from DNA (51–55). Given that Atad5-RLC represents only a minor fraction of total RFC/RLCs in *Xenopus* egg extracts, we next compared the PCNA unloading capability of all four RFC/RLC complexes. We previously established an assay that specifically detects PCNA unloading in *Xenopus* egg extracts (13). Briefly, we load human PCNA (hPCNA) with human RFC, 2–10 trimers per DNA on average, onto nicked-circular DNA immobilized on Sepharose beads, ligate the nick, wash out unbound proteins including RFC and DNA ligase, and incubate the hPCNA-DNA complex in the extracts (Fig. 2*A*). Since hPCNA can functionally replace xPCNA in NPE but is specifically detected with a monoclonal antibody that does not recognize *Xenopus* PCNA (xPCNA), this system can discriminate the unloading events of hPCNA from the possible loading events of xPCNA. In a mock-treated NPE, hPCNA was quickly unloaded from covalently closed circular DNA within 20 min (Fig. 2, *B* and *C*). Rfc3 was recovered in the DNA-bead fraction, likely reflecting the DNA-binding affinity of RFC/RLCs. In contrast, in the Rfc3-depleted NPE, where no Rfc3 binding was observed, the rate of the unloading of hPCNA was significantly retarded, suggesting that most of the unloading events in NPE depend on the Rfc3-containing complexes. Although we observed some hPCNA dissociation in Rfc3-depleted NPE, this level of PCNA dissociation likely reflects the intrinsic instability of hPCNA under our experimental condition, since a similar level of hPCNA dissociation was observed in heat-inactivated *Xenopus* egg extracts (Fig. S2). Importantly, Atad5-depletion slowed down the rate of hPCNA unloading to a similar level to that seen in Rfc3-depleted NPE (Fig. 2, *B*–*D*, see “ΔAtad5”). In contrast, Rfc1-, Ctf18-, and Rad17-depletion did not detectably reduce the rate of hPCNA unloading compared to the mock-treated NPE (Fig. 2, *B*–*D*). Rfc3 loading was significantly reduced in Rfc1-depleted NPE, suggesting that RFC has the strongest binding affinity to closed-circular DNA among all RFC/RLCs. Although these experiments do not exclude the possibility that RFC, Ctf18-RLC, and Rad17-RLC contribute to a minor fraction of hPCNA unloading, they suggest that Atad5-RLC is the major PCNA unloader in *Xenopus* egg extracts.

**Figure 2:**
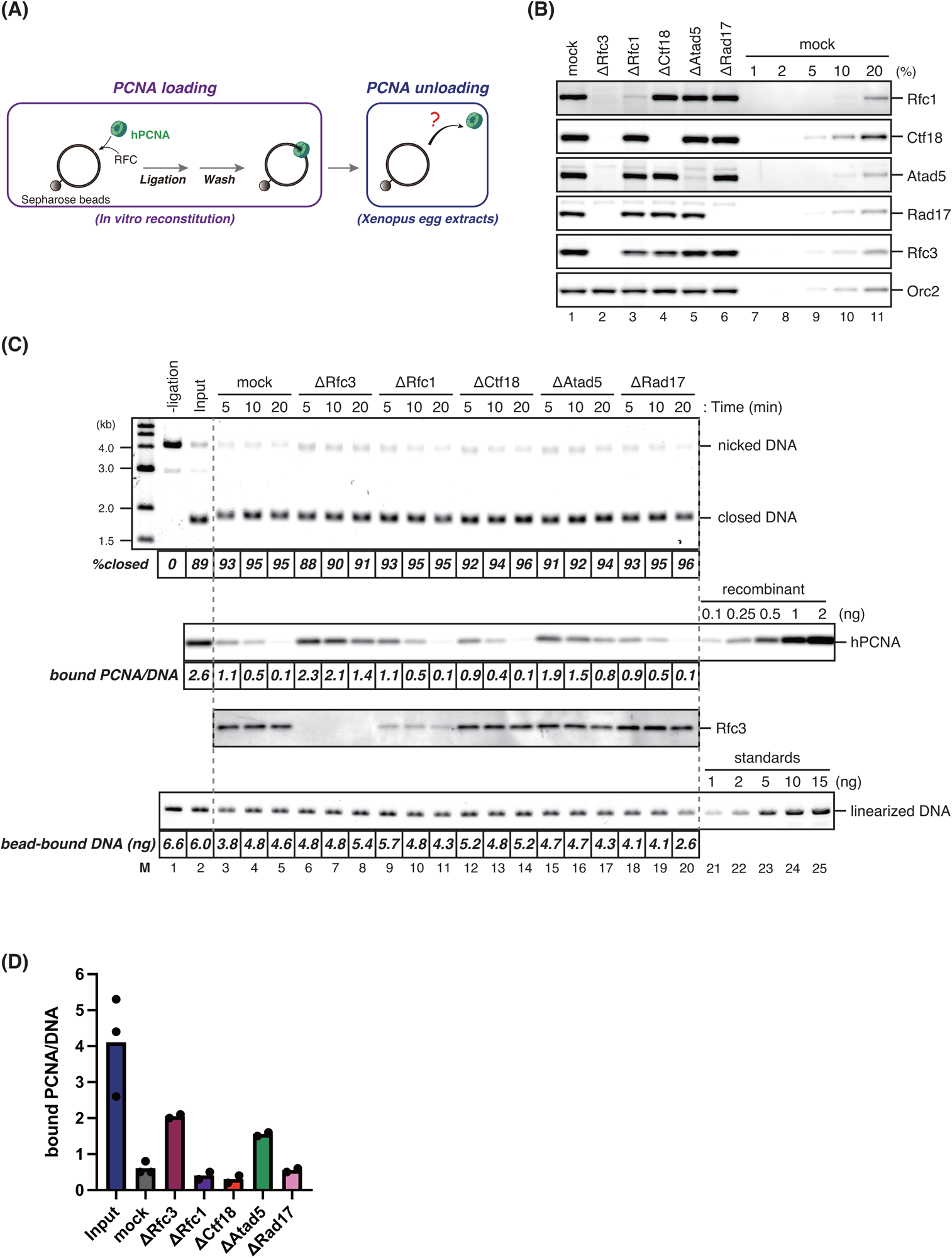
Atad5-RLC is the major PCNA unloader in *Xenopus* egg extracts. (A) A schematic diagram of the PCNA unloading assay. Singly biotinylated plasmid carrying a nick was bound to Sepharose beads and incubated with recombinant human PCNA and RFC to load PCNA on DNA. After nick ligation and extensive wash, the resulting PCNA-DNA complex was transferred into *Xenopus* egg extracts to examine PCNA unloading. (B) 0.25 μL each of mock-treated (lane 1), Rfc3-depleted (lane 2), Rfc1-depleted (lane 3), Ctf18-depleted (lane 4), Atad5-depleted (lane 5), Rad17-depleted NPE (lane 6), and a serial dilution series of mock-treated NPE (lanes 7–11) were analyzed by immunoblotting with indicated antibodies. Orc2 serves as a loading control. (C) PCNA loaded onto immobilized DNA was incubated in NPE described in (*B*). Untreated DNA separated by agarose gel electrophoresis in the presence of ethidium bromide, a quantitative immunoblot of hPCNA and Rfc3 in the bead-bound fractions, and linearized DNA (treated with XmnI) separated by agarose gel electrophoresis are presented along with the percentage of covalently-closed plasmids, the estimated numbers of DNA-bound PCNA molecules per plasmid, and the amount of bead-bound DNA. (D) The average numbers of DNA-bound PCNA per plasmid at the 10-min time points were plotted into a graph. Filled circles represent individual data from two experiments, including the one shown in (*C*).

To study further the regulation of the PCNA unloading activity of Atad5-RLC, we expressed full-length human ATAD5 with N-terminal monomeric Azami-Green (mAG) and C-terminal FLAG tags in HEK293T cells, either wild-type or a mutant form of the Walker A motif (K1138E), along with smaller RFC2–5 subunits (46), and purified the respective complexes by FLAG affinity chromatography (Fig. 3, *A* and *B*). We found that the high-speed supernatant (HSS) of *Xenopus* eggs containing both cytosolic and nuclear proteins supports efficient PCNA unloading depending on Atad5 (Fig. 3*C*, lanes 1–5), and decided to use HSS as an easy-to-make alternative to NPE. Consistent with the prediction that Atad5-RLC is the major PCNA unloader in *Xenopus* egg extracts, the wild-type ATAD5-RLC restored PCNA unloading not only in Atad5-depleted HSS but also in Rfc3-depleted HSS (Fig. 3, *C* and *D*). In contrast, ATAD5-RLC carrying the K1138E mutation was not capable of restoring PCNA unloading in Atad5-depleted HSS (Fig. 3, *E* and *F*). These results suggest that PCNA unloading by Atad5-RLC depends on its ATP-binding or hydrolysis.

**Figure 3:**
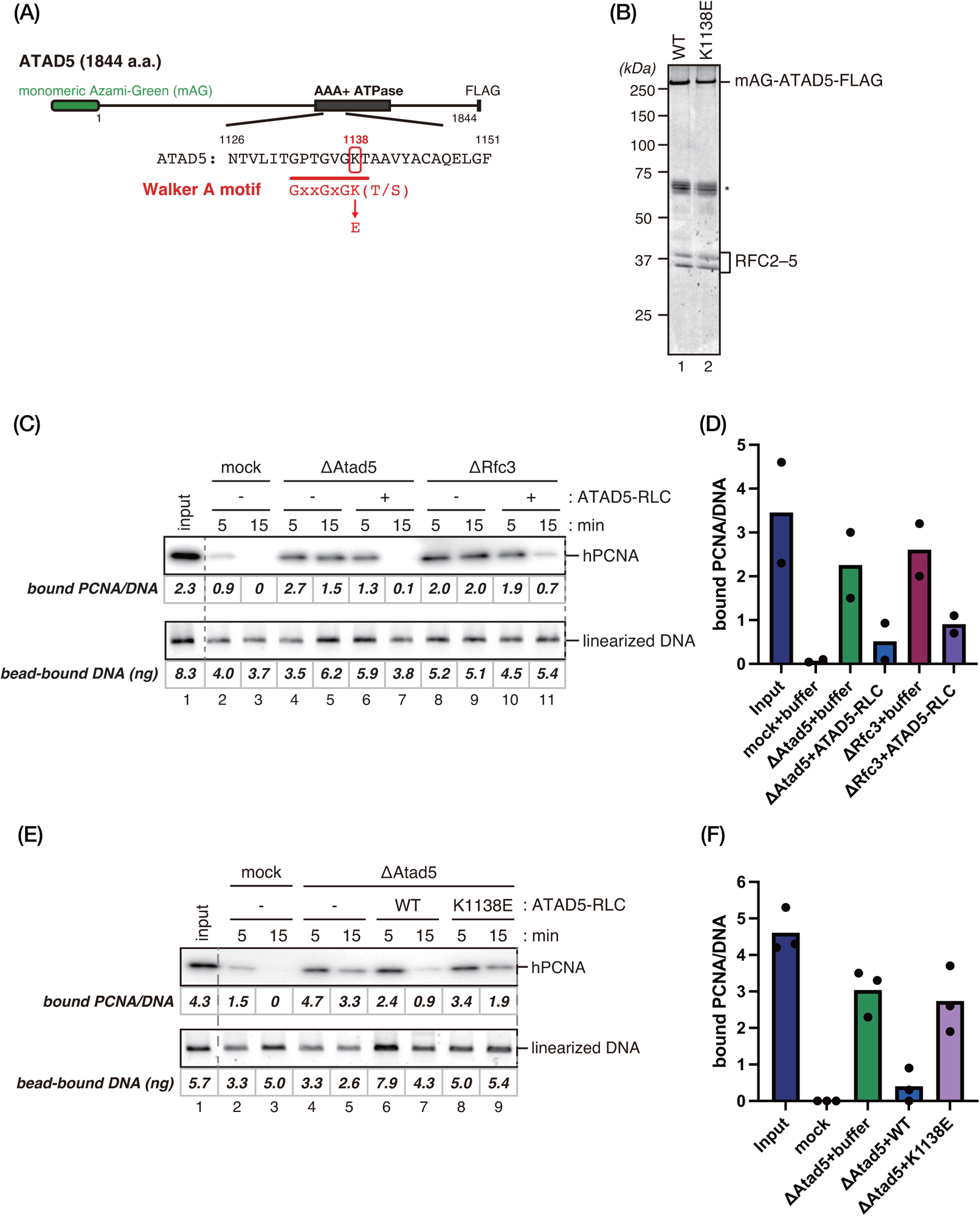
The Walker A motif of ATAD5 is required for PCNA unloading. (A) A schematic diagram of the human ATAD5 expression construct is presented. Monomeric Azami-Green (mAG) and a FLAG epitope were fused at the N- and C-termini, respectively. The Walker A mutant of ATAD5 was made by substituting the conserved lysine 1138 residue with glutamic acid. (B) 250 ng each of purified wild-type (WT) and the Walker A mutant (K1138E) ATAD5-RLCs were separated by SDS-PAGE and stained with Coomassie brilliant blue R-250. (*) indicates unrelated proteins co-purified with ATAD5-RLC. (C) The PCNA unloading assay in mock-treated HSS supplemented with buffer (lanes 2 and 3), Atad5-depleted HSS supplemented with buffer (lanes 4 and 5) or 12 nM ATAD5-RLC (lanes 6 and 7), or Rfc3-depleted HSS supplemented with buffer (lanes 8 and 9) or 12 nM ATAD5-RLC (lanes 10 and 11). An immunoblot of DNA-bound PCNA (top) and linearized DNA separated by agarose gel electrophoresis (bottom) are presented, along with the estimated numbers of PCNA molecules per DNA and the amounts of DNA. (D) The average numbers of DNA-bound PCNA per plasmid at the 15-min time points were plotted into a graph. Filled circles represent individual data from two experiments, including the one shown in (*C*). (E) The PCNA unloading assay in mock-treated HSS supplemented with buffer (lanes 2 and 3), or Atad5-depleted HSS supplemented with buffer (lanes 4 and 5), 12 nM ATAD5^WT^-RLC (lanes 6 and 7), or ATAD5^K1138E^-RLC (lanes 8 and 9). An immunoblot ofDNA-bound PCNA (top) and linearized DNA (bottom) are presented, along with the estimated numbers of PCNA molecules per DNA and the amounts of DNA. (F) The average numbers of DNA-bound PCNA per plasmid at the 15-min time points were plotted into a graph. Filled circles represent individual data from three experiments, including the one shown in (*E*).

Kubota et al. have shown that PCNA unloading occurs only after the completion of Okazaki fragment ligation in yeast (58). To gain further insight into the regulation of PCNA unloading by Atad5-RLC, we next tested whether single-strand nicks or chromatin assembly affects PCNA unloading in *Xenopus* egg extracts. First, we repeated our PCNA unloading assay in the presence of the Nb.BtsI nicking enzyme, which cleaves our DNA substrate at two loci. This enzyme efficiently converted closed-circular DNA into the open-circular form, suggesting continuous nicking in HSS, which supports efficient nick ligation (Fig. 4*A*, top panel). Despite nearly complete conversion of the plasmids into the open form, however, the rate of PCNA unloading was not detectably affected by the presence of the nicking enzyme (Fig. 4*A*, lanes 2–8, and *B*). Also, the observed unloading was largely inhibited by Atad5-depletion (Fig. 4*A*, lanes 9–14, and *B*). These results suggest that Atad5-RLC can unload PCNA regardless of the presence of single-strand nicks in *Xenopus* egg extracts.

**Figure 4:**
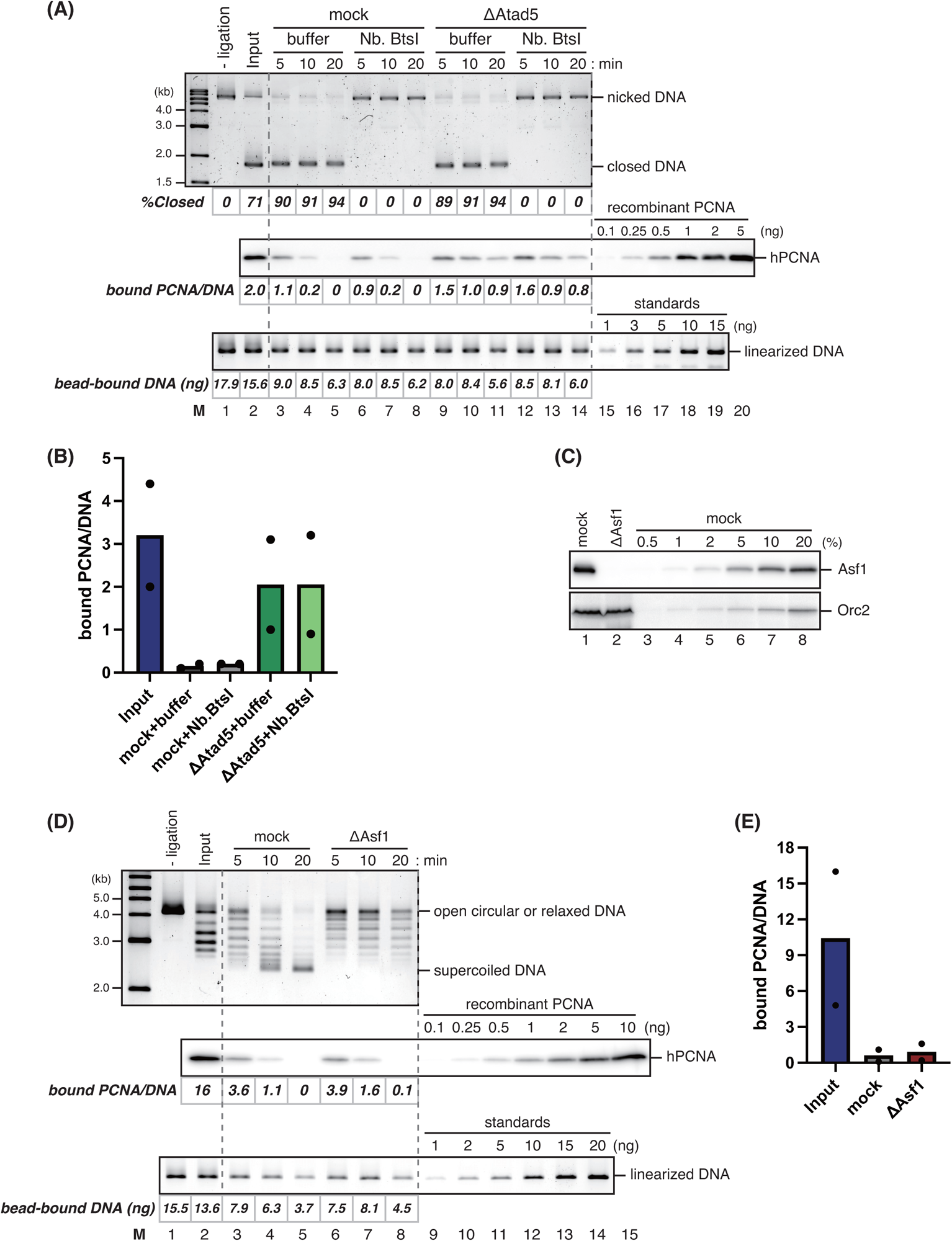
Single-strand nicks and chromatin assembly do not detectably affect PCNA unloading in *Xenopus* egg extracts. (A) The PCNA unloading assay in mock-treated HSS supplemented with buffer (lanes 3–5) or Nb.BtsI (lanes 6–8), or Atad5-depleted HSS supplemented with buffer (lanes 9–11) or Nb.BtsI (lanes 12–14), presented with the input samples before (lane 1) and after nick ligation (lane 2). Untreated DNA separated by agarose gel electrophoresis in the presence of ethidium bromide (top), a quantitative immunoblot of DNA-bound PCNA (middle), and linearized DNA separated by agarose gel electrophoresis (bottom) are shown, along with the estimated numbers of PCNA molecules per DNA and the amounts of DNA. (B) The average numbers of DNA-bound PCNA per plasmid at the 10-min time points were plotted into a graph. Filled circles represent individual data from two experiments, including the one shown in (*A*). (C) 2 μL each of mock-treated (lane 1) and Asf1-depleted HSS (lane 2) were analyzed by immunoblotting with indicated antibodies, along with a serial dilution series of mock-treated HSS (lanes 3–8). Orc2 serves as a loading control. (D) The PCNA unloading assay in HSS described in (*C*). Untreated DNA (top), a quantitative immunoblot of DNA-bound PCNA (middle), and linearized DNA (bottom) are presented, along with the estimated numbers of DNA-bound PCNA molecules per plasmid and the amounts of bead-bound DNA. (E) The average numbers of DNA-bound PCNA per plasmid at the 10-min time points were plotted into a graph. Filled circles represent individual data from two experiments, including the one shown in (*D*)

Since the completion of DNA synthesis also involves nucleosome assembly, we next tested whether nucleosome assembly affects PCNA unloading. Asf1 is the histone chaperone critical for histone deposition in both replication-coupled (CAF-1-dependent) and replication-independent (HIRA-dependent) pathways, and the depletion of Asf1 effectively inhibits nucleosome assembly in *Xenopus* egg extracts (59). Immunodepletion of Asf1 from HSS indeed prevented the supercoiling of the plasmid substrate, a readout of nucleosome deposition (Fig. 4, *C*–*E*, see “ΔAsf1”). However, it did not significantly affect the rate of PCNA unloading, suggesting that PCNA unloading occurs regardless of chromatin assembly in *Xenopus* egg extracts.

We have previously shown that the MutSα mismatch sensor complex prevents PCNA unloading to maintain MMR capability (13). Since this reaction largely depends on the PIP motif located on the N-terminus of the Msh6 subunit of MutSα, a plausible scenario is that the binding of the PIP peptide to the hydrophobic pocket on PCNA inhibits PCNA unloading. To test this possibility, we repeated the PCNA unloading assay in the presence of various PCNA-binding peptides. The PIP-peptide of p21 is one of the strongest PCNA-binding peptides, and it indeed effectively prevented PCNA unloading in HSS (Fig. 5, *A* and *B*). Both alanine mutations in the PIP consensus motif and jumbling of the peptide sequence completely inactivated the inhibitory effect of the peptide on PCNA unloading. Likewise, the PIP-peptide from MutSα impeded PCNA unloading, an effect that was neutralized by the PIP mutation and the jumbling of the peptide sequence (Fig. 5, *C* and *D*). These results suggest that PCNA unloading depends on the access of Atad5-RLC to the PIP-binding pocket on PCNA.

**Figure 5:**
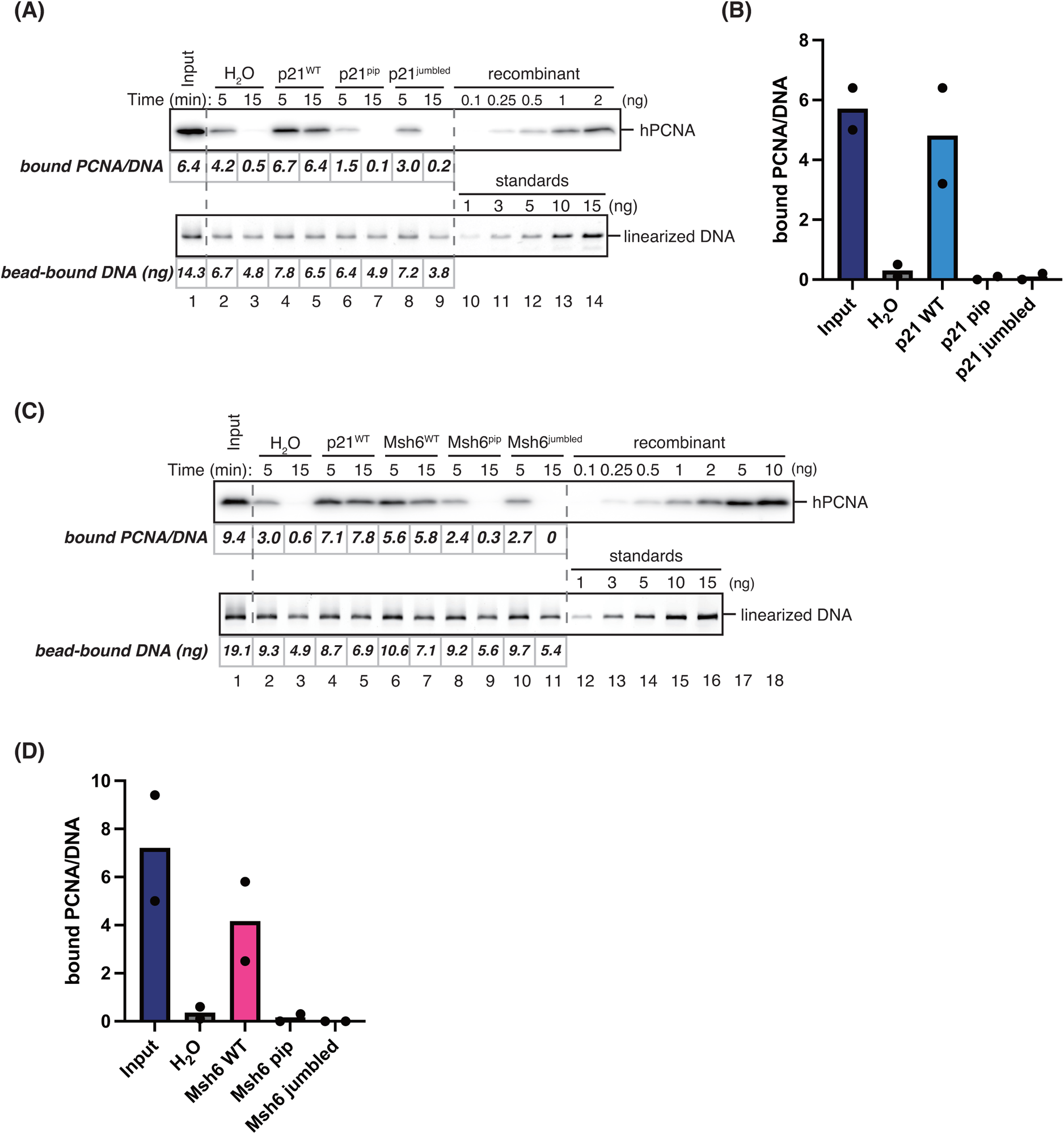
PCNA-binding peptides inhibit PCNA unloading. (A) The PCNA unloading assay in HSS supplemented with the solvent (H_2_O, lanes 2 and 3), wild-type (p21^WT^, lanes 4 and 5), PIP-mutant (p21^pip^, lanes 6 and 7), or sequence-scrambled p21 peptides (p21^jumbled^, lanes 8 and 9). A quantitative immunoblot of DNA-bound PCNA (top) and linearized DNA separated by agarose gel electrophoresis (bottom) are presented, along with the estimated numbers of PCNA molecules per DNA and the amounts of DNA. (B) The average numbers of DNA-bound PCNA per plasmid at the 15-min time points were plotted into a graph. Filled circles represent individual data from two experiments, including the one shown in (*A*). (C) The PCNA unloading assay in HSS supplemented with the solvent (H_2_O, lanes 2 and 3), wild-type p21 peptide (p21^WT^, lanes 4 and 5), wild-type (Msh6^WT^, lanes 6 and 7), PIP-mutant (Msh6^pip^, lanes 8 and 9), or sequence-scrambled Msh6 peptides (Msh6^jumbled^, lanes 10 and 11). A quantitative immunoblot of DNA-bound PCNA (top) and linearized DNA (bottom) are presented, along with the estimated numbers of PCNA molecules per DNA and the amounts of DNA. (D) The average numbers of DNA-bound PCNA per plasmid at the 15-min time points were plotted into a graph. Filled circles represent individual data from two experiments, including the one shown in (*C*).

## Discussion

By using a method that explicitly distinguishes PCNA unloading from its loading, we showed here that Atad5-RLC provides the major PCNA unloading activity in *Xenopus* egg extracts. Thus, Atad5-depletion attenuated the rate of PCNA unloading to the level comparable to Rfc3-depletion, purified ATAD5-RLC restored PCNA unloading in not only Atad5-but also Rfc3-depleted extracts, and Rfc1-, Ctf18-, or Rad17-depletion did not detectably affect the PCNA unloading kinetics (Figs. 2 and 3). Given the fact that Atad5-RLC constitutes only approximately 3% of RFC/RLCs (Fig. 1), it is conceivable that Atad5-RLC is the RFC-family clamp-loader/unloader complex that is specialized for PCNA unloading. Recent structural studies show that the Rfc1 subunit dominates the DNA binding interface over the Rfc2–5 smaller subunits (60–62), and the Rad24 subunit of Rad24-RLC directs its binding to the double-stranded region at a 5′ junction of single-stranded and double-stranded DNA in an arrangement opposite to RFC (36, 37). Therefore, it is highly likely that the largest subunits critically regulate the DNA binding specificity of RFC/RLCs, thereby restricting the situation in which each complex functions. Since almost all Rfc3 and Rfc5 are included in the RFC/RLC complexes (Figs. 1 and S1*A*), the large subunits may limit the PCNA loading/unloading activity residing in the smaller subunits.

A key regulation of PCNA unloading is to distinguish PCNA molecules that are no longer used from those in use. Kang et al. showed that replication proteins inhibit PCNA unloading by ATAD5-RLC *in vitro* (47). In good agreement with their findings, we found that PCNA-interacting peptides strongly impede PCNA unloading in *Xenopus* egg extracts (Fig. 5). Therefore, even in a physiological condition, the interaction between PCNA and PIP interactors hampers PCNA unloading, likely preventing the access of PCNA unloaders to PCNA. This model provides a reasonable framework as to how PCNA-mediated reactions are regulated: the completion of DNA synthesis, repair, and Okazaki fragment maturation liberates PCNA from its interactors, allowing the access of the PCNA unloader to DNA-bound PCNA. Conversely, providing PIP peptides around DNA-bound PCNA would be an effective strategy to prevent PCNA unloading. PCNA is a strand-discrimination marker for eukaryotic MMR (12), and ongoing MMR inhibits PCNA unloading to maintain the strand-discrimination capability (13). The MMR system likely does so by loading multiple MutSα onto a mismatch site and providing a high concentration of the PIP peptides, each of which is located at the N-terminus of a long and flexible linker on the Msh6 subunit (63).

## Experimental procedures

### Preparation of *Xenopus* egg extracts

*Xenopus laevis* was purchased from Kato-S-Science (Chiba, Japan) and handled according to the animal experimental regulations at Kyushu University. NPE and HSS were prepared as described previously (64, 65).

### Cloning

Symbols of human and *Xenopus* genes and proteins conformed to the nomenclature guidelines of HGNC (https://www.genenames.org/about/guidelines/) and Xenbase (https://www.xenbase.org/entry/static/gene/geneNomenclature.jsp).

Cloning of the *Xenopus laevis rfc3* gene was performed as follows: The *rfc3* gene was amplified from *Xenopus* egg cDNA by two-step PCR using primers listed in Supporting Table S1 and cloned into pDONR201 (Thermo Fisher Scientific, MA, USA) by the Gateway BP reaction, resulting in pDONR-xRFC3. For protein expression in *Escherichia coli*, the *rfc3* gene on pDONR-xRFC3 was transferred into pET-HSD, an in-house Gateway destination vector carrying a T7 promoter and an N-terminal His_6_-tag, by the Gateway LR reaction.

Cloning of the human *ATAD5* gene was performed as follows: The *ATAD5* gene was amplified from the HeLa cDNA library with a FLAG-tag sequence fused to the C-terminus of ATAD5 by two-step PCR using primers listed in Supporting Table S1, digested with BamHI (New England Biolabs, MA, USA, #R3136) and SbfI (New England Biolabs, #R3642), and ligated with a BamHI/SbfI-digested vector fragment that had been amplified by PCR from pCSII-EF-mAG-TEV-6His-Claspin-3Flag (66) using primers listed in Supporting Table S1, resulting in pCSII-EF-mAG-hATAD5-FLAG. The Walker A motif mutant of hATAD5 (hATAD5-K1138E) was constructed by two-step PCR using primers listed in Supporting Table S1. The amplified fragment was digested with BamHI and HpaI (New England Biolabs, #R0105) and inserted between the BamHI and HpaI sites in pCSII-EF-mAG-hATAD5-FLAG, resulting in pCSII-EF-mAG-hATAD5-K1138E-FLAG.

### Protein expression and purification

Expression and purification of wild-type and the Walker A mutant (K1138E) of ATAD5-RLC, human PCNA, and human RFC were carried out as described previously (13, 67).

Expression and purification of the His_6_-tagged *Xenopus* Rfc3 protein were performed as follows: Recombinant protein expression was induced in *E.coli* BL21-CodonPlus (DE3)-RIPL cells transformed with pET-HSD-xRFC3 by the addition of 0.5 mM Isopropyl β-D-1-thiogalactopyranoside (IPTG) in Terrific broth for 2 h at 37°C. Cells were harvested, lysed with 1 mg/mL lysozyme, sonicated in buffer S (50 mM Na-phosphate pH 8.0, 300 mM NaCl, 5% glycerol, 0.1% Triton-X100, 2.5 mM 2-mercaptoethanol, 1 mM phenylmethylsulfonyl fluoride [PMSF]), and centrifuged at 20,400 ×*g* for 10 min. Inclusion bodies containing the Rfc3 protein were resuspended in buffer S, sonicated, and centrifuged again at 20,400 ×*g* for 5 min, and the procedure was repeated twice. The Rfc3 protein was extracted from purified inclusion bodies with Laemmli’s SDS sample buffer.

### Immunological methods

Production and usage of antibodies against *Xenopus* Rfc3 were described previously (13). The mouse monoclonal antibody against human PCNA (MBL International Corporation, #MH-12-3) was commercially available. The rabbit antiserum against *Xenopus* Orc2 was a kind gift from Dr. Johannes Walter. The rabbit Ctf18, Rad17, Rfc5, Ctf8, and Asf1 antibodies were raised against peptides NH_2_-CINEEFGENDSEILENDDNA-COOH, corresponding to residues 269–287 of Ctf18, NH_2_-CAAQAIMEDEELKIEEYDSD-COOH, corresponding to residues 269–287 of Rad17, NH_2_-CEHVVKEERVDISPDGMK-COOH, corresponding to residues 182–198 of Rfc5, NH_2_-CKIIFKTRPKPIITNVPKKV-COOH, corresponding to residues 103–121 of Ctf8, NH_2_-APSKGLAAALNTLPENSMDC-COOH, corresponding to residues 180–199 of Asf1, respectively. All antibodies except anti-Ctf8 and anti-Asf1 were affinity-purified using corresponding antigens. Polyclonal antibodies against *Xenopus* Atad5 and Rfc1 were raised in rabbit against the following polypeptides: Atad5: N-terminal 105 amino acids of Atad5 with a 6-histidine tag at the C-terminus (MVGILAMSASLEEYGCQPCKKSRKDEEAPIKTITNYFS PVSKNTEKVLSSPRSNNIADYFKQNSPINEKKQTSKAENAAIQQDTPQAAVADSSAAS GKPSKCRKR); Rfc1: N-terminal 100 amino acids of Rfc1 with a 6-histidine tag at the C-terminus (MDIRNFFGVKPVAKKHGTEKTDIKEKKKSPEAKKKPKDSKVKSPSS DDSLKGMNVKKKKRIIYDSDAEEESPPVKKAKKPSEKSPPLPRPHKIRKPDPVV).

For immunoblotting, the Orc2 antiserum was used at a dilution of 1:10,000, and the Rfc1, Atad5, Ctf8, and Asf1 antisera were used at 1:5,000 dilutions. For immunoblotting of Ctf18, Rad17, Rfc3, and Rfc5, affinity-purified antibodies were used at 0.5 μg/mL. Alexa fluor 647 conjugated Goat anti-rabbit IgG (H+L) antibodies (Jackson ImmunoResearch, #111-605-144), Goat anti-mouse IgG (H+L) antibodies (Jackson ImmunoResearch, #115-035-146), and Monoclonal Mouse anti-rabbit IgG specific to the light chain (Jackson ImmunoResearch, #211-602-171) were used at 1:10,000 dilutions as the secondary antibodies.

For immunodepletion of Rfc1, Atad5, or Asf1, 3 vol of an antiserum was bound to 1 vol of recombinant Protein A Sepharose Fast Flow (PAS; Cytiva, #17127902) at 4°C overnight. For Ctf18, Rad17, or Rfc3 depletion, 5 μg of purified IgG was bound to 1 μL PAS. To deplete extracts, 0.2 vol of the antibody-coupled PAS beads were incubated with 1 vol of NPE or HSS at 4°C for 1 h, and the procedure was repeated twice. In most cases, we depleted 20∼60 μL of extracts for an experiment. For quadruple depletion of Rfc1, Ctf18, Atad5, and Rad17 from NPE, antibody-coupled beads were prepared separately as described above, 2 μL each of IgG beads were combined, and 8 μL of the combined IgG beads were incubated in 10 μL of NPE that had been diluted to 50 μL (5-fold) with Egg lysis buffer salts (ELB-salts: 10 mM Hepes-KOH pH 7.7, 2.5 mM MgCl_2_, 50 mM KCl) at 4°C for 1 h, and the procedure was repeated twice.

### Quantification of RFC/RLCs in *Xenopus* egg extracts

The concentration of RFC and other RLCs was estimated from the amount of Rfc3 co-precipitated with the largest subunits. For IP of Rfc1 or Atad5, 1.5 vol of an antiserum was bound to 1 vol of PAS, and for IP of Rfc3, Ctf18, or Rad17, 15 μg of affinity-purified antibodies were bound to 1 μL PAS at 4°C overnight. NPE was diluted 10-fold with ELB-salts and clarified by centrifugation at 20,400 ×*g* for 10 min to remove insoluble debris. 10 μL each of antibody-coupled beads were incubated with 100 μL of diluted NPE at 4°C for 1h, washed three times with ELB-salts, and bound proteins were eluted with 40 μL of Laemmli’s SDS sample buffer. IP samples were separated by SDS-PAGE, alongside a dilution series of recombinant Rfc3, transferred onto polyvinylidene fluoride membranes, and probed with antibodies against Rfc3 and other subunits. The amounts of Rfc3 in the IP fractions were calculated using the dilution series of recombinant Rfc3 as a standard.

### Preparation of substrates for the *in vitro* PCNA-loading assay

*In vitro* synthesis of plasmids carrying a site-specific biotin modification was performed as described previously (13). Briefly, an oligonucleotide carrying a biotin-dT modification was annealed on a 3-kb single-stranded phagemid DNA, and the complementary DNA strand was synthesized by in-house-purified T7 DNA polymerase. After ligation of nicks by T4 DNA ligase (Nippon gene, Tokyo, Japan, #311-00404), covalently closed DNA molecules were purified by cesium chloride density gradient ultra-centrifugation. To make a PCNA loading site, a locus-specific nick was introduced by Nt.BbvCI (New England Biolabs, #R0632). The DNA was bound onto Sepharose beads at a ratio of 100 ng per 1 μL, following the procedure described previously (68).

### *In vitro* PCNA-loading assay

The *in vitro* PCNA-loading assay was performed as described previously, with some modifications (13). Briefly, DNA beads were washed three times with mHBS buffer (10 mM Hepes-NaOH pH 7.5, 0.05% Tween-20, 10 mM MgCl_2_, 200 μM ethylenediaminetetraacetic acid [EDTA], 150 mM NaCl) and incubated in 2 vol of mHBS containing 50 mM phosphocreatine (PC), 25 μg/mL creatine phosphokinase (CPK), 2 mM adenosine triphosphate (ATP), 400 μM dithiothreitol (DTT), 145 ng/μL hPCNA, and 2.2 ng/μL RFC at 32°C for 15 min. The PCNA-DNA complex was washed three times with mHBS and incubated in 4 vol of ligation buffer (50 mM Tris-HCl pH 7.9, 10 mM, MgCl_2_, 20 mM DTT, 1 mM ATP) containing 25 units/μL T4 DNA ligase at 32°C for 5 min, washed three times with mHBS, once with ELB-salts containing 1M KCl, and then once with ELB-salts.

### The PCNA unloading assay in *Xenopus* egg extracts

*Xenopus* egg extracts (HSS and NPE) were supplemented with 2 mM ATP, 20 mM PC, 5 μg/mL CPK, and pre-incubated at 22°C for 5 min. The PCNA-DNA complex was incubated in the extract at the concentration of 20 ng/μL with respect to immobilized DNA at 22°C. At appropriate time points, the mixture was quickly diluted with 200 μL of ELB-salts containing 0.2% Triton X-100, overlayed onto 200 μL of ELB-salts containing 0.5 M Sucrose and centrifuged at 12,700 ×*g* for 1 min at 4°C in a TMS-21 swinging bucket rotor (Tomy Seiko, Tokyo, Japan, #0621650333). The beads were washed once with 200 μL of ELB-salts and resuspended in an appropriate volume of ELB-salts buffer. DNA beads were then split into two aliquots to quantify the amount of bead-bound DNA and DNA-loaded PCNA. To quantify DNA, an aliquot of the beads was mixed with 100 μL of stop buffer (1% sodium dodecyl sulfate [SDS] and 20 mM EDTA) and treated with 50 μg/mL Proteinase K (Nacalai Tesque, Kyoto, Japan, #29442-14) at 37°C for 1 h. DNA was then purified by phenol/chloroform extraction and ethanol precipitation and resuspended in TE buffer (10 mM Tris-HCl pH 7.4, 1 mM EDTA) containing 10 μg/mL RNase A. DNA samples were digested with XmnI (New England Biolabs, #R0194) and analyzed by 0.8% agarose gel electrophoresis followed by staining with SYBR Gold nucleic acid gel stain (Thermo Fisher Scientific, #S11494). Fluorescent signals were detected using the Amersham Typhoon scanner 5 system (Cytiva, MA, USA) and processed using the ImageQuant TL software (Cytiva). To evaluate the PCNA amount, another aliquot of the beads was mixed with Laemmli’s SDS sample buffer, and DNA-bound proteins were analyzed by SDS-PAGE followed by immunoblotting. The p21 and Msh6 peptides (p21 wild-type, NH_2_-MSKQKTLFSFFTKSPPVSSS-COOH; p21 pip, NH_2_-MSKAKTLFSAATKSPPVSSS-COOH; p21 jumbled, NH_2_-SVSSFTPKLQSFKSPTSFKM-COOH; Msh6 wild-type, NH_2_-KRRQTSMTDFYHSKRRLIFS-COOH; Msh6 pip, NH_2_-KRRATSMTDAAHSKRRLIFS-COOH; Msh6 jumbled, NH_2_-QDKTRYFHRTMSRSKSIRLF-COOH) (69) were synthesized by Eurofins Genomics (Luxemburg, Luxemburg), and added to HSS at the final concentration of 1 mg/mL. To introduce nicks on plasmids, Nb.BtsI (New England Biolabs, #R0707) was added to HSS at the concentration of 0.5 units/μL. For heat-inactivation, HSS was boiled at 95°C for 5 min, centrifuged at 20,400 ×*g* for 10 min at 4°C, and the supernatant was recovered as heat-inactivated HSS.

### Data availability

All data from this study are included in the manuscript and supporting information. Unprocessed raw data and DNA sequences underlying this article will be shared on reasonable request to the corresponding authors.

## Supporting information

This article contains supporting information. Supporting figures include co-depletion of the small subunits with the large subunits in NPE, the recombinant protein used for quantification (Fig. S1), and the stability of DNA-bound hPCNA in heat-inactivated *Xenopus* egg extracts (Fig. S2). Supporting Table S1 summarizes the oligonucleotide primers used in this study.

## Acknowledgments

We thank Dr. Johannes Walter for antibodies and Dr. Hisao Masai for sharing plasmids for expression in HEK293T cells. We also thank Dr. Eiji Ohashi for critical reading and members of the Takahashi lab for helpful discussions.

## Author contributions

Yoshitaka Kawasoe: Conceptualization, Funding acquisition, Formal analysis, Investigation (most experiments), Methodology, Writing—original draft. Sakiko Shimokawa: Investigation (Fig. 3), Methodology, Resources. Peter J. Gillespie: Resources, Writing—review & editing. J. Julian Blow: Funding acquisition, Resources, Writing—review & editing. Toshiki Tsurimoto: Conceptualization, Methodology, Resources. Tatsuro S. Takahashi: Conceptualization, Funding acquisition, Supervision, Methodology, Writing—review & editing.

## ORCID

Yoshitaka Kawasoe: 0000-0003-0925-2004

Peter J. Gillespie: 0000-0002-8812-5267

J. Julian Blow: 0000-0002-9524-5849

Toshiki Tsurimoto: 0000-0001-7597-2216

Tatsuro S. Takahashi: 0000-0002-1947-7680

## Abbreviations

PCNA: proliferating cell nuclear antigen
hPCNA: human PCNA
xPCNA: *Xenopus* PCNA
RFC: Replication factor C
Ctf18: Chromosome transmission fidelity 18
Atad5: ATPase family AAA domain containing 5
Elg1: enhanced level of genome instability 1
RLC: RFC-like complex
MMR: mismatch repair
PIP: PCNA-interacting peptide
9–1–1: Rad9–Hus1–Rad1
NPE: nucleoplasmic extract
HSS: high-speed supernatant
IP: Immunoprecipitation
PAS: Protein A Sepharose
IgG: immunoglobulin G
ELB-salts: Egg lysis buffer salts
mHBS: modified HEPES-buffered Saline
PC: phosphocreatine
CPK: creatine phosphokinase

## References

1. Burgers, P. M. J., and Kunkel, T. A. (2017) Eukaryotic DNA Replication Fork Annu Rev Biochem 86, 417–438 10.1146/annurev-biochem-061516-044709

2. Kunkel, T. A., and Erie, D. A. (2015) Eukaryotic Mismatch Repair in Relation to DNA Replication Annual Review of Genetics 49, 291–313 10.1146/annurev-genet-112414-054722

3. Boehm, E. M., Gildenberg, M. S., and Washington, M. T. (2016) The Many Roles of PCNA in Eukaryotic DNA Replication Enzymes 39, 231–254 10.1016/bs.enz.2016.03.003

4. Choe, K. N., and Moldovan, G. L. (2017) Forging Ahead through Darkness: PCNA, Still the Principal Conductor at the Replication Fork Molecular Cell 65, 380–392 10.1016/j.molcel.2016.12.020

5. Jonsson, Z. O., Hindges, R., and Hubscher, U. (1998) Regulation of DNA replication and repair proteins through interaction with the front side of proliferating cell nuclear antigen EMBO J 17, 2412–2425 10.1093/emboj/17.8.2412

6. Warbrick, E. (1998) PCNA binding through a conserved motif BioEssays 20, 195–199 10.1002/(sici)1521-1878(199803)20:3<195::Aid-bies2>3.0.Co;2-r

7. Flores-Rozas, H., Clark, D., and Kolodner, R. D. (2000) Proliferating cell nuclear antigen and Msh2p-Msh6p interact to form an active mispair recognition complex Nat Genet 26, 375–378 10.1038/81708

8. Kleczkowska, H. E., Marra, G., Lettieri, T., and Jiricny, J. (2001) hMSH3 and hMSH6 interact with PCNA and colocalize with it to replication foci Genes Dev 15, 724–736 10.1101/gad.191201

9. Goellner, E. M., Smith, C. E., Campbell, C. S., Hombauer, H., Desai, A., Putnam, C. D. et al. (2014) PCNA and Msh2-Msh6 activate an Mlh1-Pms1 endonuclease pathway required for Exo1-independent mismatch repair Mol Cell 55, 291–304 10.1016/j.molcel.2014.04.034

10. Hombauer, H., Campbell, C. S., Smith, C. E., Desai, A., and Kolodner, R. D. (2011) Visualization of eukaryotic DNA mismatch repair reveals distinct recognition and repair intermediates Cell 147, 1040–1053 10.1016/j.cell.2011.10.025

11. Kadyrov, F. A., Dzantiev, L., Constantin, N., and Modrich, P. (2006) Endonucleolytic function of MutLalpha in human mismatch repair Cell 126, 297–308 10.1016/j.cell.2006.05.039

12. Pluciennik, A., Dzantiev, L., Iyer, R. R., Constantin, N., Kadyrov, F. A., and Modrich, P. (2010) PCNA function in the activation and strand direction of MutLalpha endonuclease in mismatch repair Proc Natl Acad Sci U S A 107, 16066–16071 10.1073/pnas.1010662107

13. Kawasoe, Y., Tsurimoto, T., Nakagawa, T., Masukata, H., and Takahashi, T. S. (2016) MutSalpha maintains the mismatch repair capability by inhibiting PCNA unloading Elife 5, 10.7554/eLife.15155

14. Pillon, M. C., Babu, V. M., Randall, J. R., Cai, J., Simmons, L. A., Sutton, M. D., et al. (2015) The sliding clamp tethers the endonuclease domain of MutL to DNA Nucleic Acids Res 43, 10746–10759 10.1093/nar/gkv918

15. Lee, S. D., and Alani, E. (2006) Analysis of interactions between mismatch repair initiation factors and the replication processivity factor PCNA J Mol Biol 355, 175–184 10.1016/j.jmb.2005.10.059

16. Kadyrov, F. A., Holmes, S. F., Arana, M. E., Lukianova, O. A., O’Donnell, M., Kunkel, T. A., et al. (2007) Saccharomyces cerevisiae MutLalpha is a mismatch repair endonuclease J Biol Chem 282, 37181–37190 10.1074/jbc.M707617200

17. Umar, A., Buermeyer, A. B., Simon, J. A., Thomas, D. C., Clark, A. B., Liskay, R. M. et al. (1996) Requirement for PCNA in DNA mismatch repair at a step preceding DNA resynthesis Cell 87, 65–73, https://www.ncbi.nlm.nih.gov/pubmed/8858149

18. Iyer, R. R., Pluciennik, A., Genschel, J., Tsai, M. S., Beese, L. S., and Modrich, P. (2010) MutLalpha and proliferating cell nuclear antigen share binding sites on MutSbeta J Biol Chem 285, 11730–11739 10.1074/jbc.M110.104125

19. Ohashi, E., and Tsurimoto, T. (2017) Functions of Multiple Clamp and Clamp-Loader Complexes in Eukaryotic DNA Replication Adv Exp Med Biol 1042, 135–162 10.1007/978-981-10-6955-0_7

20. Arbel, M., Choudhary, K., Tfilin, O., and Kupiec, M. (2021) PCNA Loaders and Unloaders-One Ring That Rules Them All Genes (Basel) 12, 10.3390/genes12111812

21. Lee, K. Y., and Park, S. H. (2020) Eukaryotic clamp loaders and unloaders in the maintenance of genome stability Exp Mol Med 52, 1948–1958 10.1038/s12276-020-00533-3

22. Shiomi, Y., and Nishitani, H. (2017) Control of Genome Integrity by RFC Complexes; Conductors of PCNA Loading onto and Unloading from Chromatin during DNA Replication Genes (Basel) 8, 10.3390/genes8020052

23. Fairman, M., Prelich, G., Tsurimoto, T., and Stillman, B. (1988) Identification of cellular components required for SV40 DNA replication in vitro Biochim Biophys Acta 951, 382–387 10.1016/0167-4781(88)90110-8

24. Bowman, G. D., O’Donnell, M., and Kuriyan, J. (2004) Structural analysis of a eukaryotic sliding DNA clamp-clamp loader complex Nature 429, 724–730 10.1038/nature02585

25. Gomes, X. V., Schmidt, S. L., and Burgers, P. M. (2001) ATP utilization by yeast replication factor C. II. Multiple stepwise ATP binding events are required to load proliferating cell nuclear antigen onto primed DNA J Biol Chem 276, 34776–34783 10.1074/jbc.M011743200

26. Miyata, T., Oyama, T., Mayanagi, K., Ishino, S., Ishino, Y., and Morikawa, K. (2004) The clamp-loading complex for processive DNA replication Nat Struct Mol Biol 11, 632–636 10.1038/nsmb788

27. Zhuang, Z., Yoder, B. L., Burgers, P. M., and Benkovic, S. J. (2006) The structure of a ring-opened proliferating cell nuclear antigen-replication factor C complex revealed by fluorescence energy transfer Proc Natl Acad Sci U S A 103, 2546–2551 10.1073/pnas.0511263103

28. Johnson, A., Yao, N. Y., Bowman, G. D., Kuriyan, J., and O’Donnell, M. (2006) The replication factor C clamp loader requires arginine finger sensors to drive DNA binding and proliferating cell nuclear antigen loading J Biol Chem 281, 35531–35543 10.1074/jbc.M606090200

29. Hanna, J. S., Kroll, E. S., Lundblad, V., and Spencer, F. A. (2001) Saccharomyces cerevisiae CTF18 and CTF4 are required for sister chromatid cohesion Mol Cell Biol 21, 3144–3158 10.1128/MCB.21.9.3144-3158.2001

30. Mayer, M. L., Gygi, S. P., Aebersold, R., and Hieter, P. (2001) Identification of RFC(Ctf18p, Ctf8p, Dcc1p): an alternative RFC complex required for sister chromatid cohesion in S. cerevisiae Mol Cell 7, 959–970 10.1016/s1097-2765(01)00254-4

31. Kanellis, P., Agyei, R., and Durocher, D. (2003) Elg1 forms an alternative PCNA-interacting RFC complex required to maintain genome stability Curr Biol 13, 1583–1595 10.1016/s0960-9822(03)00578-5

32. Bellaoui, M., Chang, M., Ou, J., Xu, H., Boone, C., and Brown, G. W. (2003) Elg1 forms an alternative RFC complex important for DNA replication and genome integrity EMBO J 22, 4304–4313 10.1093/emboj/cdg406

33. Ben-Aroya, S., Koren, A., Liefshitz, B., Steinlauf, R., and Kupiec, M. (2003) ELG1, a yeast gene required for genome stability, forms a complex related to replication factor C Proc Natl Acad Sci U S A 100, 9906–9911 10.1073/pnas.1633757100

34. Green, C. M., Erdjument-Bromage, H., Tempst, P., and Lowndes, N. F. (2000) A novel Rad24 checkpoint protein complex closely related to replication factor C Curr Biol 10, 39–42 10.1016/s0960-9822(99)00263-8

35. Merkle, C. J., Karnitz, L. M., Henry-Sanchez, J. T., and Chen, J. (2003) Cloning and characterization of hCTF18, hCTF8, and hDCC1. Human homologs of a Saccharomyces cerevisiae complex involved in sister chromatid cohesion establishment J Biol Chem 278, 30051–30056 10.1074/jbc.M211591200

36. Castaneda, J. C., Schrecker, M., Remus, D., and Hite, R. K. (2022) Mechanisms of loading and release of the 9-1-1 checkpoint clamp Nat Struct Mol Biol 29, 369–375 10.1038/s41594-022-00741-7

37. Zheng, F., Georgescu, R. E., Yao, N. Y., O’Donnell, M. E., and Li, H. (2022) DNA is loaded through the 9-1-1 DNA checkpoint clamp in the opposite direction of the PCNA clamp Nat Struct Mol Biol 29, 376–385 10.1038/s41594-022-00742-6

38. Day, M., Oliver, A. W., and Pearl, L. H. (2022) Structure of the human RAD17-RFC clamp loader and 9-1-1 checkpoint clamp bound to a dsDNA-ssDNA junction Nucleic Acids Res 50, 8279–8289 10.1093/nar/gkac588

39. Murakami, T., Takano, R., Takeo, S., Taniguchi, R., Ogawa, K., Ohashi, E., et al. (2010) Stable interaction between the human proliferating cell nuclear antigen loader complex Ctf18-replication factor C (RFC) and DNA polymerase epsilon is mediated by the cohesion-specific subunits, Ctf18, Dcc1, and Ctf8 J Biol Chem 285, 34608–34615 10.1074/jbc.M110.166710

40. Grabarczyk, D. B., Silkenat, S., and Kisker, C. (2018) Structural Basis for the Recruitment of Ctf18-RFC to the Replisome Structure 26, 137–144 e133 10.1016/j.str.2017.11.004

41. Fujisawa, R., Ohashi, E., Hirota, K., and Tsurimoto, T. (2017) Human CTF18-RFC clamp-loader complexed with non-synthesising DNA polymerase ε efficiently loads the PCNA sliding clamp Nucleic Acids Research 45, 4550–4563 10.1093/nar/gkx096

42. Liu, H. W., Bouchoux, C., Panarotto, M., Kakui, Y., Patel, H., and Uhlmann, F. (2020) Division of Labor between PCNA Loaders in DNA Replication and Sister Chromatid Cohesion Establishment Mol Cell 78, 725–738 e724 10.1016/j.molcel.2020.03.017

43. Baris, Y., Taylor, M. R. G., Aria, V., and Yeeles, J. T. P. (2022) Fast and efficient DNA replication with purified human proteins Nature 606, 204–210 10.1038/s41586-022-04759-1

44. Kubota, T., Nishimura, K., Kanemaki, M. T., and Donaldson, A. D. (2013) The Elg1 replication factor C-like complex functions in PCNA unloading during DNA replication Mol Cell 50, 273–280 10.1016/j.molcel.2013.02.012

45. Shiomi, Y., and Nishitani, H. (2013) Alternative replication factor C protein, Elg1, maintains chromosome stability by regulating PCNA levels on chromatin Genes Cells 18, 946–959 10.1111/gtc.12087

46. Lee, K. Y., Fu, H., Aladjem, M. I., and Myung, K. (2013) ATAD5 regulates the lifespan of DNA replication factories by modulating PCNA level on the chromatin J Cell Biol 200, 31–44 10.1083/jcb.201206084

47. Kang, M. S., Ryu, E., Lee, S. W., Park, J., Ha, N. Y., Ra, J. S., et al. (2019) Regulation of PCNA cycling on replicating DNA by RFC and RFC-like complexes Nat Commun 10, 2420 10.1038/s41467-019-10376-w

48. Bell, D. W., Sikdar, N., Lee, K. Y., Price, J. C., Chatterjee, R., Park, H. D., et al. (2011) Predisposition to cancer caused by genetic and functional defects of mammalian Atad5 PLoS Genet 7, e1002245 10.1371/journal.pgen.1002245

49. Johnson, C., Gali, V. K., Takahashi, T. S., and Kubota, T. (2016) PCNA Retention on DNA into G2/M Phase Causes Genome Instability in Cells Lacking Elg1 Cell Rep 16, 684–695 10.1016/j.celrep.2016.06.030

50. Paul Solomon Devakumar, L. J., Gaubitz, C., Lundblad, V., Kelch, B. A., and Kubota, T. (2019) Effective mismatch repair depends on timely control of PCNA retention on DNA by the Elg1 complex Nucleic Acids Res 47, 6826–6841 10.1093/nar/gkz441

51. Yao, N. Y., Johnson, A., Bowman, G. D., Kuriyan, J., and O’Donnell, M. (2006) Mechanism of proliferating cell nuclear antigen clamp opening by replication factor C J Biol Chem 281, 17528–17539 10.1074/jbc.M601273200

52. Cai, J., Gibbs, E., Uhlmann, F., Phillips, B., Yao, N., O’Donnell, M., et al. (1997) A complex consisting of human replication factor C p40, p37, and p36 subunits is a DNA-dependent ATPase and an intermediate in the assembly of the holoenzyme J Biol Chem 272, 18974–18981, https://www.ncbi.nlm.nih.gov/pubmed/9228079

53. Yao, N., Turner, J., Kelman, Z., Stukenberg, P. T., Dean, F., Shechter, D., et al. (1996) Clamp loading, unloading and intrinsic stability of the PCNA, beta and gp45 sliding clamps of human, E. coli and T4 replicases Genes Cells 1, 101–113, https://www.ncbi.nlm.nih.gov/pubmed/9078370

54. Shibahara, K., and Stillman, B. (1999) Replication-dependent marking of DNA by PCNA facilitates CAF-1-coupled inheritance of chromatin Cell 96, 575–585, https://www.ncbi.nlm.nih.gov/pubmed/10052459

55. Bylund, G. O., and Burgers, P. M. (2005) Replication protein A-directed unloading of PCNA by the Ctf18 cohesion establishment complex Mol Cell Biol 25, 5445–5455 10.1128/MCB.25.13.5445-5455.2005

56. Walter, J., Sun, L., and Newport, J. (1998) Regulated chromosomal DNA replication in the absence of a nucleus Mol Cell 1, 519–529 10.1016/s1097-2765(00)80052-0

57. Hoogenboom, W. S., Klein Douwel, D., and Knipscheer, P. (2017) Xenopus egg extract: A powerful tool to study genome maintenance mechanisms Dev Biol 428, 300–309 10.1016/j.ydbio.2017.03.033

58. Kubota, T., Katou, Y., Nakato, R., Shirahige, K., and Donaldson, A. D. (2015) Replication-Coupled PCNA Unloading by the Elg1 Complex Occurs Genome-wide and Requires Okazaki Fragment Ligation Cell Rep 12, 774–787 10.1016/j.celrep.2015.06.066

59. Shintomi, K., Inoue, F., Watanabe, H., Ohsumi, K., Ohsugi, M., and Hirano, T. (2017) Mitotic chromosome assembly despite nucleosome depletion in Xenopus egg extracts Science 356, 1284–1287 10.1126/science.aam9702

60. Gaubitz, C., Liu, X., Pajak, J., Stone, N. P., Hayes, J. A., Demo, G., et al. (2022) Cryo-EM structures reveal high-resolution mechanism of a DNA polymerase sliding clamp loader Elife 11, 10.7554/eLife.74175

61. Schrecker, M., Castaneda, J. C., Devbhandari, S., Kumar, C., Remus, D., and Hite, R. K. (2022) Multistep loading of a DNA sliding clamp onto DNA by replication factor C Elife 11, 10.7554/eLife.78253

62. Zheng, F., Georgescu, R., Yao, N. Y., Li, H., and O’Donnell, M. E. (2022) Cryo-EM structures reveal that RFC recognizes both the 3’- and 5’-DNA ends to load PCNA onto gaps for DNA repair Elife 11, 10.7554/eLife.77469

63. Shell, S. S., Putnam, C. D., and Kolodner, R. D. (2007) The N terminus of Saccharomyces cerevisiae Msh6 is an unstructured tether to PCNA Mol Cell 26, 565–578 10.1016/j.molcel.2007.04.024

64. Terui, R., Nagao, K., Kawasoe, Y., Taki, K., Higashi, T. L., Tanaka, S. et al. (2018) Nucleosomes around a mismatched base pair are excluded via an Msh2-dependent reaction with the aid of SNF2 family ATPase Smarcad1 Genes Dev 32, 806–821 10.1101/gad.310995.117

65. Sparks, J., and Walter, J. C. (2018) Extracts for Analysis of DNA Replication in a Nucleus-Free System Cold Spring Harb Protoc 10.1101/pdb.prot097154

66. Uno, S., and Masai, H. (2011) Efficient expression and purification of human replication fork-stabilizing factor, Claspin, from mammalian cells: DNA-binding activity and novel protein interactions Genes Cells 16, 842–856 10.1111/j.1365-2443.2011.01535.x

67. Miyashita, R., Nishiyama, A., Qin, W., Chiba, Y., Kori, S., Kato, N., et al. (2023) The termination of UHRF1-dependent PAF15 ubiquitin signaling is regulated by USP7 and ATAD5 Elife 12, 10.7554/eLife.79013

68. Higashi, T. L., Ikeda, M., Tanaka, H., Nakagawa, T., Bando, M., Shirahige, K. et al. (2012) The prereplication complex recruits XEco2 to chromatin to promote cohesin acetylation in Xenopus egg extracts Curr Biol 22, 977–988 10.1016/j.cub.2012.04.013

69. Mattock, H., Jares, P., Zheleva, D. I., Lane, D. P., Warbrick, E., and Blow, J. J. (2001) Use of peptides from p21 (Waf1/Cip1) to investigate PCNA function in Xenopus egg extracts Exp Cell Res 265, 242–251 10.1006/excr.2001.5181

