## Supporting materials for "The Atad5 RFC-like complex is the major unloader of proliferating cell nuclear antigen in *Xenopus* egg extracts"

Figure S1

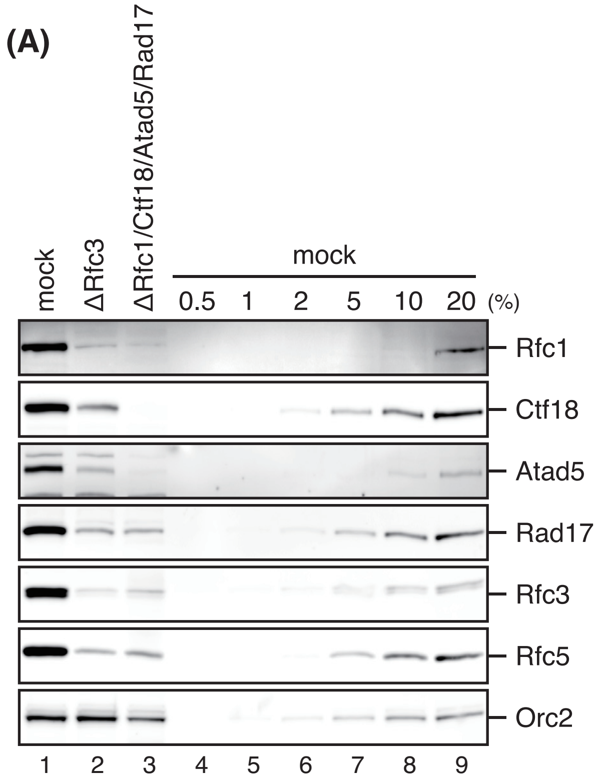
(A)
mock
ΔRfc3
ΔRfc1/Ctf18/Atad5/Rad17
mock
0.5
1
2
5
10
20
(%)
Rfc1
Ctf18
Atad5
Rad17
Rfc3
Rfc5
Orc2
1
2
3
4
5
6
7
8
9

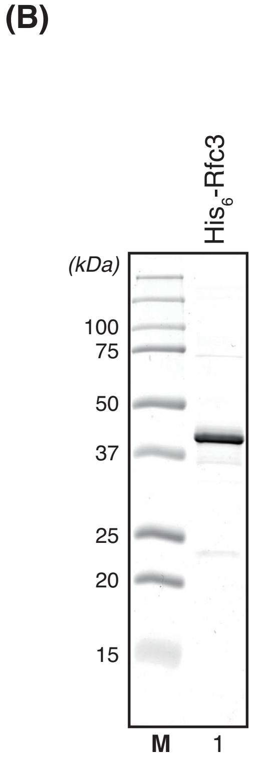
(B)
His6-Rfc3
(kDa)
100
75
50
37
25
20
15
M
1

**Supporting Figure S1: Most of Rfc3 and Rfc5 are included in RFC and RLCs in *Xenopus* egg extracts**

(*A*) 0.25 µL each of mock-treated (lane 1), Rfc3-depleted (lane 2), and Rfc1/Ctf18/Atad5/Rad17-depleted NPE (lane 3) were analyzed by immunoblotting with indicated antibodies alongside a serial dilution series of mock-treated NPE (lanes 4–9). Orc2 serves as a loading control. Quadruple depletion of Rfc1, Ctf18, Atad5, and Rad17 removed more than 90% of Rfc3 and Rfc5, suggesting that most of the small subunits in *Xenopus* egg extracts are included in RFC and RLCs.

(*B*) 2.3 µg of recombinant $His_6$-Rfc3 purified from *E. coli* was separated by SDS-PAGE and stained with Coomassie brilliant blue R-250.

**Figure S2**

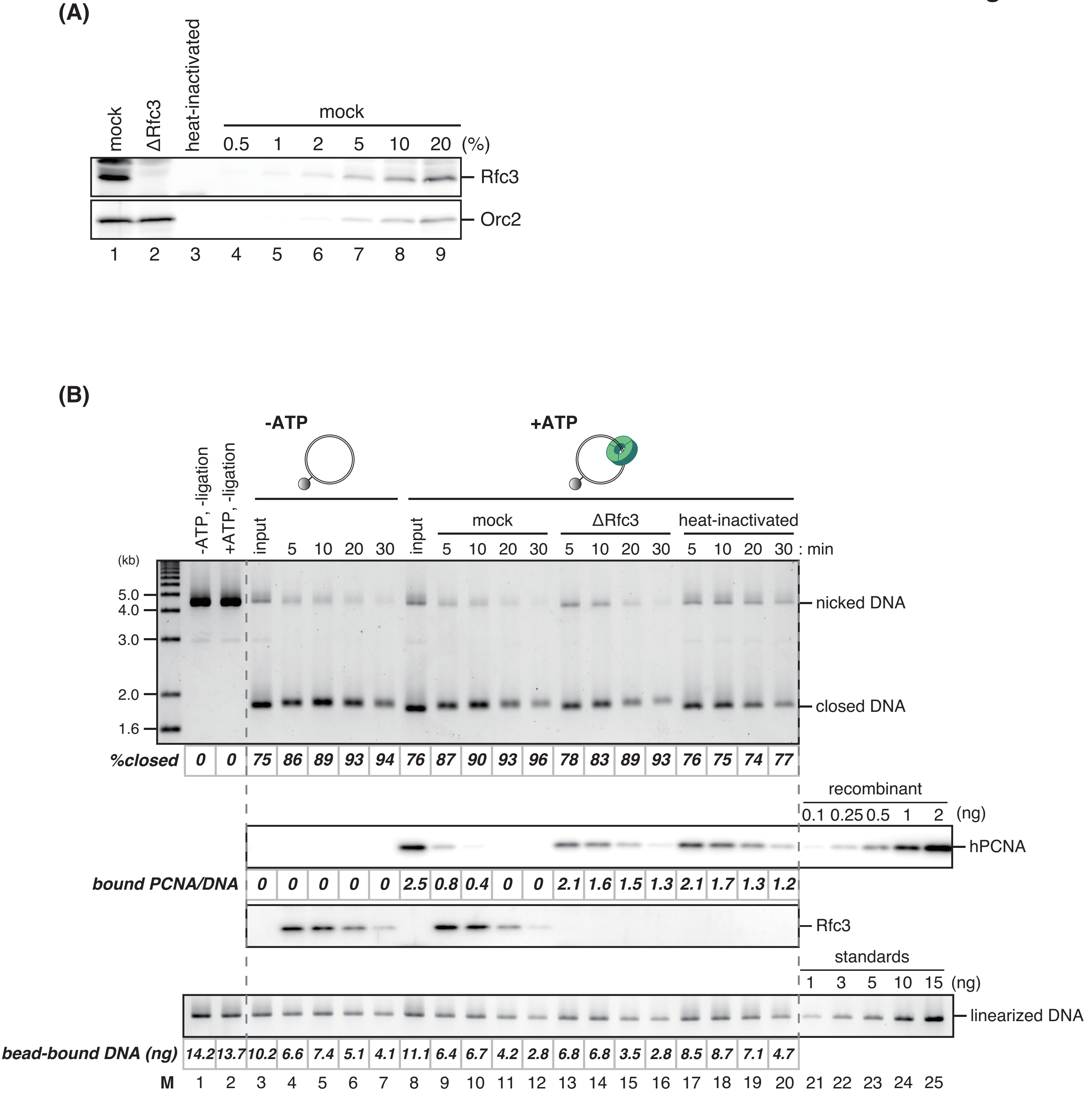
(A)
mock
ΔRfc3
heat-inactivated
mock
0.5 1 2 5 10 20 (%)
Rfc3
Orc2
1 2 3 4 5 6 7 8 9
(B)
-ATP
+ATP
-ATP, -ligation
+ATP, -ligation
input
5 10 20 30
input
mock
ΔRfc3
heat-inactivated
5 10 20 30 5 10 20 30 5 10 20 30 : min
(kb)
5.0
4.0
3.0
2.0
1.6
nicked DNA
closed DNA
%closed
0 0 75 86 89 93 94 76 87 90 93 96 78 83 89 93 76 75 74 77
recombinant
0.1 0.25 0.5 1 2 (ng)
hPCNA
bound PCNA/DNA
0 0 0 0 0 2.5 0.8 0.4 0 0 2.1 1.6 1.5 1.3 2.1 1.7 1.3 1.2
Rfc3
standards
1 3 5 10 15 (ng)
linearized DNA
bead-bound DNA (ng)
14.2 13.7 10.2 6.6 7.4 5.1 4.1 11.1 6.4 6.7 4.2 2.8 6.8 6.8 3.5 2.8 8.5 8.7 7.1 4.7
M 1 2 3 4 5 6 7 8 9 10 11 12 13 14 15 16 17 18 19 20 21 22 23 24 25

**Supporting Figure S2: Spontaneous dissociation of PCNA from DNA in heat-inactivated *Xenopus* egg extracts**

(*A*) 2 µL each of mock-treated (lane 1), Rfc3-depleted (lane 2), and heat-inactivated HSS (lane 3), from which heat-denatured insoluble proteins had been removed, were analyzed by immunoblotting with indicated antibodies along with a serial dilution series of mock-treated HSS (lanes 4–9). Orc2 serves as a loading control.

(*B*) The *in vitro* PCNA loading assay in the absence (lanes 1 and 3–7) or the presence of ATP (lanes 2 and 8–20) in mock-treated (lanes 4–7 and 9–12), Rfc3-depleted (lanes 13–16) or heat-inactivated HSS (lanes 17–20). Untreated DNA separated by agarose gel electrophoresis in the presence of ethidium bromide (top), a quantitative immunoblot of hPCNA and Rfc3 in the bead-bound fractions (middle), and linearized DNA separated by agarose gel electrophoresis (bottom) are presented along with the percentage of covalently-closed plasmids, the estimated numbers of DNA-bound PCNA molecules per plasmid, and the amounts of DNA. PCNA loaded onto immobilized DNA was dissociated in Rfc3-depleted and heat-inactivated HSS with similar kinetics, suggesting that the observed PCNA dissociation in Rfc3-depleted extracts reflects the spontaneous dissociation of PCNA from DNA.

**Supproting Table S1: oligonucleotides used in this study**

| Name | Sequence (5'-3') | Application | Genes |
|---|---|---|---|
| 1116 | AAAGCAGGCTCCACCATGAGTTTGTGGGTTGATAAGTATCGACC | Cloning (Forward) | *Xenopus rfc3* with a partial Gateway attB sequence |
| 1117 | ACAAGAAAGCTGGGTCTCAAAACATCATCGCTTCTAGCCCATCC | Cloning (Reverse) | *Xenopus rfc3* with a partial Gateway attB sequence |
| 344 | GGGGACAAGTTTGTACAAAAAAGCAGGCTCCAC | Cloning (Forward) | Gateway attB |
| 345 | GGGGACCACTTTGTACAAGAAAGCTGGGTC | Cloning (Reverse) | Gateway attB |
| ATAD5-1F-BamHI | GGAAGGATCCATGGTGGGGGTCCTGGCCATGGCGGC | Cloning (Forward) and Mutagenesis (Forward, Fragment 1) | *human ATAD5* |
| ATAD5-5535R | TTAAGGGAAGTCAGCTGCCAAAGTATTTACAGTCTC | Cloning (Reverse) | *human ATAD5* |
| ATAD5-5532R-FL-SbfI | GGAACCTGCAGGTTACTTGTCATCGTCATCCTTGTAGTCTCGAGGGAAGTCAGCTGCCAAAGTATTTAC | Fusion of a FLAG-tag and a SbfI site to ATAD5 (Reverse) | *human ATAD5* with a FLAG-tag sequence and a SbfI site |
| pCSII-EF-MCS-F | GGAACCTGCAGGGCGGCCAACATCGAGGGATCAAGCTTATCGAT | Cloning (Forward) | pCSII-EF plasmid backbone |
| pCSII-EF-mAG-R | GGAAGGATCCATGGTGGTGATGGTGGTGGC | Cloning (Reverse) | pCSII-EF plasmid backbone with mAG, TEV cleavage site, and $His_6$ sequences |
| ATAD5-K1138E-R | GCAGCAGTTTCTCCCACTCCTGTTGGCCCTG | Mutagenesis (Reverse, Fragment 1) | *human ATAD5* |
| ATAD5-K1138E-F | GGAGTGGGAGAAACTGCTGCAGTGTATGCTTG | Mutagenesis (Forward, Fragment 2) | *human ATAD5* |
| ATAD5-4163R-HpaI | GCAGTTAACAAGGTTACAAAG | Mutagenesis (Reverse, Fragment 2) | *human ATAD5* |
